# Revolutionising notifiable avian disease diagnostics: validation of direct swab testing for Avian Influenza and Newcastle disease using the CENOS platform

**DOI:** 10.64898/2026.08.11.744268

**Authors:** David Edge, James Turton, Adeola Adebo, Omeryasin Tuzaktepe, Brad Fraser, Craig Ross, Joe James, Jacob Terrey, Nelson Nazareth, Scott M Reid, Ashley C Banyard

**Affiliations:** BG RESEARCH, 5 The Business Centre, Harvard Way, Kimbolton, Cambridgeshire, PE28 0NJ, United Kingdom; BIOGENE, 6 The Business Centre, Harvard Way, Kimbolton, Cambridgeshire, PE28 0NJ, United Kingdom; Influenza and Avian Virology workgroup, Animal and Plant Health Agency, Weybridge, United Kingdom; WOAH/FAO International reference laboratory for avian influenza, swine influenza and Newcastle disease virus, Animal and Plant Health Agency, Weybridge, United Kingdom; School of Life Sciences, University of Sussex. Falmer, East Sussex

**Keywords:** High pathogenicity avian influenza virus (HPAIV), notifiable avian disease (NAD), Newcastle disease virus (NDV), frontline diagnostics, molecular testing, point-of-care testing

## Abstract

Existing molecular diagnostic approaches for notifiable avian diseases (NADs) involve a suite of PCR assays that enable both generic detection, and where positive, subtyping of both avian influenza virus (AIV) and Newcastle disease virus (NDV). Novel rapid and direct diagnostic assays for the detection of AIV and NDV were developed and evaluated using unprocessed cloacal (C) and oropharyngeal (OP) poultry swab material. Both assays employ a closed tube direct real-time reverse transcription polymerase chain reaction (RRT-PCR) approach in which viral lysis is achieved by heat treatment and a dedicated PCR compatible buffer, followed by detection using a RRT-PCR approach. Primer and probe sets were designed using globally circulating AIV and NDV sequences collected over the preceding five years, rather than region-specific sequence datasets, so that the assays detect all circulating genotypes. Analytical performance assessment demonstrated that both assays were highly sensitive and specific, successfully detecting all unextracted target antigens without cross reactivity to a panel of other common poultry pathogens. For each assay, viral lysis and amplification were achieved directly from samples at single digit genome copy numbers. Furthermore, low levels of viral RNA could be reliably detected in the presence of C and OP matrix material, providing proof-of-concept for direct detection of these economically significant avian pathogens in a field setting. Additional use case scenarios, including pooled sample screening and combined C/OP testing from individual birds, were also explored. These findings establish a foundation for ongoing studies incorporating paired-sample testing against validated laboratory reference assays.

## Short Communication

Notifiable avian diseases (NAD) are a significant obstacle to sustainable agriculture globally and have profound impacts on both wild and captive bird populations. The two globally defined NADs are avian influenza virus (AIV) and Newcastle disease virus (NDV), and together represent the most economically important poultry diseases. Both NADs cause a spectrum of clinical signs in infected birds, with host response to infection and viral diversity meaning that no clinical disease signs are pathognomonic for infection and as such suspicion against a broad range of clinical disease signs must be investigated, utilising highly sensitive and specific diagnostic tools.

AIVs constitute a complex group of viruses that can cause a range of disease outcomes from asymptomatic infection to severe disease, sometimes with very high morbidity and mortality in some bird species. AIVs circulate continuously in wild bird populations, but with the ability to spread to commercially reared birds and with the potential for spread to other taxonomic groups. AIVs are subtyped according to surface glycoproteins the haemagglutinin (HA) and the neuraminidase (NA) gene. Currently, 17 different HA proteins and 9 different NA proteins have been described in wild birds or poultry (Karakus et al., 2024; Krammer et al., 2018) (ref). Although most AIVs have a reservoir in wild waterfowl, numerous AIV subtypes circulate in poultry (e.g., H5N1, H9N2) causing significant impacts to the sector. Infection of poultry with avian influenza viruses is further categorised into notifiable and non-notifiable virus subtypes. The criteria for notifiable avian influenza subtypes are defined, alongside high pathogenicity (HPAI) and low pathogenicity (LPAI) outcomes, by the World Organisation for Animal Health (WOAH) (WOAH, 2021). Globally, the H5 and H7 AIV subtypes are classified as notifiable regardless of whether they have a HP or LP infection outcome in poultry.

The current AIV H5N1 panzootic is caused by a clade 2.3.4.4b H5N1 virus, a descendent of the A/goose/Guangdong/1/1996 (Gs/Gd) lineage that was first identified in China (Krammer et al., 2018). Since then, through both genetic shift (reassortment of genetic material) and drift (slow mutation of key residues over time) the HPAI A(H5) Gs/GD lineage viruses have spread to almost all corners of the globe, representing a serious global threat for the poultry industry, wild bird species, and representing a zoonotic risk. Since October 2020, the clade2.3.4.4b H5N1 HPAIV has caused consecutive epidemic waves across Europe and has spread globally (Fusaro et al., 2024).

Further to poultry risk, AIVs are often associated with zoonotic threat alongside their impact upon avian species. The clade 2.3.4.4b H5N1 panzootic has led to spill over of HPAIV into both terrestrial (Banyard et al., 2025; Burrough et al., 2024; Mostafa et al., 2024) and marine mammal (Kuiken et al., 2026; Sooksawasdi Na Ayudhya et al., 2025) species as well as into farmed mammalian species (Martins et al., 2025). Through spread and circulation in cattle, zoonotic risk had increased although despite numerous opportunities for human infection, cases reported in humans have been limited. Indeed, whilst the clade 2.3.4.4b H5N1 virus has dominated globally, human cases, limited to Cambodia and the surrounding region, have been caused by an alternative Gs/GD clade (Chin et al., 2026).

Newcastle disease (ND) is caused by virulent strains of *Avian paramyxovirus* type 1 (APMV-1). These viruses are negative sense, single-stranded, non-segmented, enveloped RNA virus belonging to the *Paramyxoviridae* family of viruses. APMV-1s circulate globally in avian hosts, often in the absence of clinical disease. Classification of APMV-1 is undertaken by analysis of the full fusion (F)-gene sequence (Dimitrov et al., 2019) with currently two classes of APMV-1 being defined. Like the HP/LP classification of notifiable AIVs, clinical and genetic criteria are used to define when APMV-1s are denoted as NDV in poultry (World Organisation for Animal Health (WOAH), 2021) (WOAH manual). The major determinant of virulence in APMV-1 is the presence of a multi-basic cleavage site (CS) and a phenylalanine at position 117 (World Organisation for Animal Health (WOAH), 2021). Although officially considered a zoonotic agent, infections with NDV in humans and mammals are very rare (Brown et al., 2026) and range from conjunctivitis to a fatal encephalitis (Abolnik and Hayes, 2025).

In Great Britain (GB), suspicion of NAD is currently investigated by official veterinarians and if negation in the field is not possible, samples are submitted to the National Reference Laboratory (NRL) for diagnostic testing. As both AI and ND are classified as notifiable and exotic, live virus containing samples must be worked within Specified Animal Pathogen Order (SAPO) categorised facilities at SAPO level 4. Further, because AIV can be zoonotic the samples also have to be handled within Advisory Committee for Dangerous Pathogens (ACDP; <u>House of Commons - Innovation, Universities, Science and Skills - Written Evidence</u>) Level 3 facilities. Finally, as pandemic AIVs are categorised as schedule 5 pathogens, facilities must also meet those criteria where live virus is manipulated.

Rapidity of diagnostic evaluation for NADs is critical wherever there is suspicion. Novel tools that can help expedite disease confirmation are required. Many laboratory-based methods for the detection of these pathogens have been described. For AIVs, this often requires both a generic screening PCR (Nagy et al., 2021; Spackman et al., 2002) followed by a subtyping PCR approach (Monne et al., 2008; Slomka et al., 2007) to define the presence of notifiable H5 and H7 subtypes. In some instances, pathotyping PCR assays are available to define HP versus LP strains (James et al., 2022). In the case of suspicion of ND, again, generic PCR screening is undertaken (Sutton et al., 2019; Wise et al., 2004) followed by pathotyping PCR or Sanger sequencing to determine the sequence across the cleavage site (Aldous et al., 2003; Fuller et al., 2009). However, more rapid, simpler methods would support better outbreak monitoring and surveillance screening. Whilst several in field tests have been developed, they typically have sensitivities that are several orders of magnitude lower than laboratory based molecular tests. For example, in-field applications focussed on ELISA-based antigen detection are of limited utility unless being deployed within an acute infection setting (Lin et al., 2025). Certainly, because of the spectrum of disease potential that NADs have in different species, sensitivity is critical to the diagnostic pipeline. Another important factor impacting upon assay sensitivity is sample matrix. The tolerance of assays for swab samples used in NAD diagnostic testing is thought to be a fundamental limitation whereby the enzymes used in these assays are unable to detect low titres of genomic RNA when compared to the gold-standard RT-PCR assays. This reduces their suitability for application in reactive settings as confidence in outputs is not high.

Time to diagnosis is critical when considering NADs. Standard molecular diagnostic tests are based on nucleic acid extraction processes prior to amplification, adding time to generation of diagnostic results through the requirement for complex infrastructure and expert users, and so in practice is limited to laboratory settings. To reduce laboratory and staff training requirements, a method based on simply adding the oropharyngeal (OP) or cloacal (C) swabs from birds directly into the reaction, without prior requirement for any extraction or processing has been developed to expedite and simplify the diagnostic pathway.

To enable this approach validation of a ‘direct from swab’ testing pipeline for surveillance and front-line diagnosis of these two economically important diseases of poultry has been developed. The approach is an enhancement on previous studies demonstrating extraction free molecular diagnosis for Peste des Petits Ruminants (PPR) virus and Foot and Mouth Disease (FMD) virus (Edge et al., 2022). Raw material was utilised as samples in an extraction free process to rapidly diagnose the presence or absence of these disease agents in unprocessed goat and cattle nasal swab eluates, respectively (Edge et al., 2022). Here, we have demonstrated that the same basic methodology can be applied to the detection of NAD from poultry oropharyngeal or cloacal swabs.

It was envisaged that C swabs would be more inhibitory to the process than the OP swabs due to the presence of faecal material. To undertake the evaluation, commercially available AIV (BG-KIT-AIV01) and NDV (BG-KIT-NDV01) kits from BioGene (Kimbolton, UK) were developed. These reagents were designed based on an alignment of all occurring AIV and class II NDV sequences observed globally during the preceding five years. As such, these tests can detect >99.5% of circulating strains while relying on a multiple primer approach to ensure full coverage of genotypes and lineages as required. The kit utilises a direct detection methodology **(Figure 1)**. The chemistry supporting the active reagent contains a mixture of thermostable enzymes capable of performing both RNA and DNA directed DNA polymerase activity in a closed tube RT-PCR based detection system. Any infectious virus is lysed during the initial denaturation step, and multiple cycles of reverse transcription are then made possible by the thermostable enzymes. This approach enables high assay sensitivity, since there is no extraction or concentration during the process. In a normal RT-PCR, the RT enzyme is thermally denatured during the initial PCR denaturation, when both the RNA target and the enzyme will be present for multiple cycles until the RNA has degraded. This cyclical RT (CRT) process ensures that the assays can detect down to single figure target copies in the presence of crude sample (**Figure 1**).

**Fig 1.**
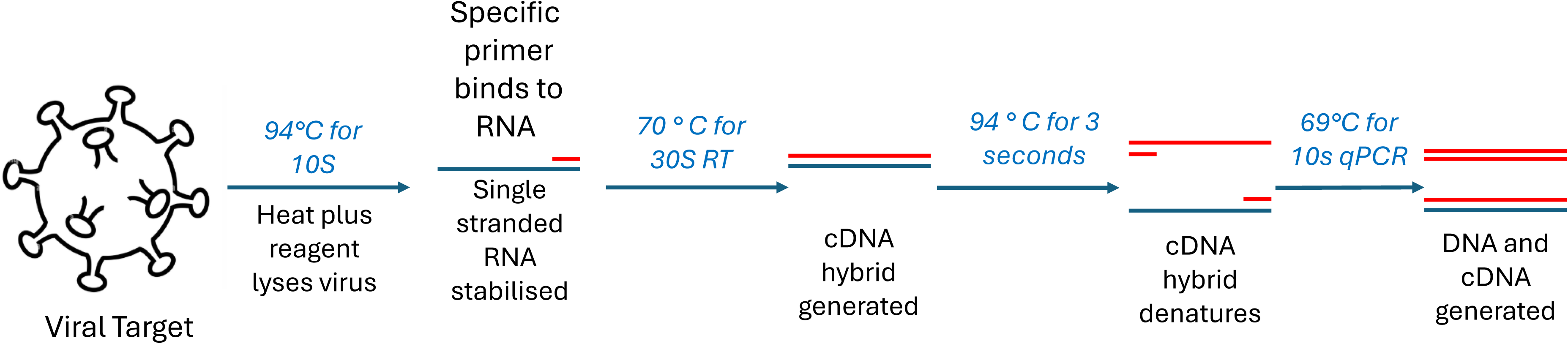
Diagrammatic representation of the detection methodology. The reagents lyse the viruses in the crude sample, releasing nucleic acids and stabilizing them. Detection is performed via high temperature Cyclical RT-PCR, CRT-PCR, to maximise sensitivity during extraction free detection from crude samples. Specific detection relies on the use of hydrolysis probes to perform qPCR.

The performance of the novel assay was compared to the standard frontline RT-PCR assay to assess sensitivity. This was initially undertaken using *in vitro* transcribed (IVT) RNA of representative AIV and NDV strains as quantified standards (**Table 1**). The AIV and NDV detection assays, individually, had a lower limit of detection (LLOD) than the frontline PCR assays with less than ten targets per reaction being detected and with reverse transcription being successfully performed across a wide range of temperatures (**Supplementary Figure 1**). From this evaluation, the novel assays were able to detect spiked IVT RNA to as low as 6.25 IVT targets for AIV and 16.25 IVT targets for NDV.

**Table 1:** Summary of the virus strains used in the study.

| Sample type | Virus / subtype | Strain | Abbreviation |
| --- | --- | --- | --- |
| IVT RNA | NDV | Chicken/Nigeria/Vom/VRD216/73/2013 | CN |
| IVT RNA | NDV | PPMV-1/Belgium/05-03936-8/2005 | BP |
| IVT RNA | NDV | Duck/CN/JX/79C2/2016 | DC |
| Antigen | NDV | APMV-1/chicken/N. Ireland/Ulster/67 | UL |
| Antigen | NDV | APMV-1/Pigeon/UK/AV0167/2021 | PP |
| Antigen | NDV | APMV-1/chicken/USA/Lasota/1946 | LS |
| IVT RNA | AIV | A/Falco_peregrinus/Belgium/03518_0004/2023 | FP |
| IVT RNA | AIV | Black-headed_gull/Finland/7826_23VIR6803-11/2023 | BG |
| IVT RNA | AIV | A/chicken/England/062653/2023 | IC |
| Inactivated antigen | AIV H5N1 | A/Turkey/England/031170/25 | TE |
| Inactivated antigen | AIV H5N1 | A/Chicken/Wales/053969/21 | CW |
| Inactivated antigen | AIV H5N5 | A/Mute_swan/Croatia/102/2016 | SC |
| Inactivated antigen | AIV H7N3 | A/Chicken/England/40054/06 | CE |
| Inactivated antigen | AIV H7N9 | A/Anhui/1/13 | CA |

The next step was to evaluate the novel assay using sample material in an unextracted format from different sample matrices. This was critical as for the novel assay, the aim was to eliminate the need for viral RNA (vRNA) extraction and undertake the amplification in the presence of significant amounts of host material captured as part of the sampling process. Further, demonstration that live viral material is rendered inactive as part of the assay is critical for the assay to enable rapid manipulation in lower containment settings. To determine that both AIV and the causative agent of NDV were successfully lysed and rendered amplifiable by the reagent, we generated standard curves from dilutions of infectious viral material covering a range of virus subtypes (**Figure 2**). This demonstrated that both AIV subtypes and NDV lineages could be amplified using the new assay and that the method was semi-quantitative. Further, for AIV, we demonstrated that the novel methodology was able to detect a broad range of H5N1 genotypes, including those currently representing a significant global threat to animal and human health. To undertake this work, reactions were spiked with 7-log serial dilutions of five AIV and three NDV unextracted viral preparations. All samples were detected across the full standard curve (**Figure 2)**.

**Figure 2.**
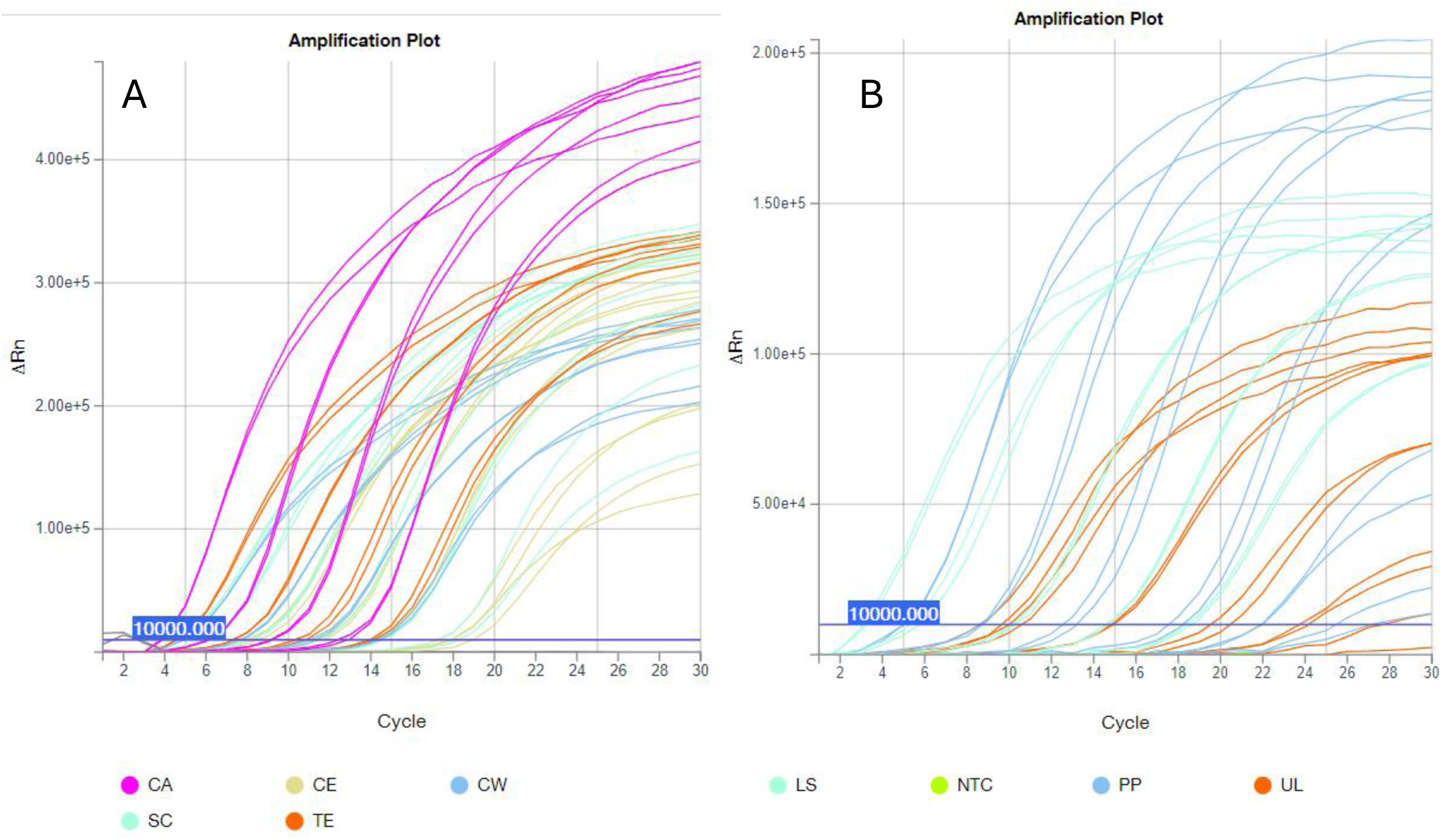
The AIV assay (A) and NDV assay (B), serial dilutions of unextracted and inactivated viral antigens to determine that the assays detected multiple genotypes. For the AIV assay (A) five isolates were chosen (Table 1) across a 7-fold serial dilution and for NDV (B) 3 isolates were treated likewise. No template controls were included and were negative for both assays out to cycle 50.

Samples matrix testing was then undertaken by adding swab eluate, both from cloacal and oropharyngeal swabs, into reaction mixes containing IVT RNA templates near the LLOD. The aim was to determine if the presence of the crude samples, which contain known inhibitors such as faecal and oral exudate, would negatively impact the sensitivity of either assay. With the AIV assay, across 60 cloacal and 25 oropharyngeal swabs there was maximally a one PCR cycle offset in Cq value but no difference in the LLOD in the presence of the swab eluate material (**Figure 3**).

**Figure 3.**
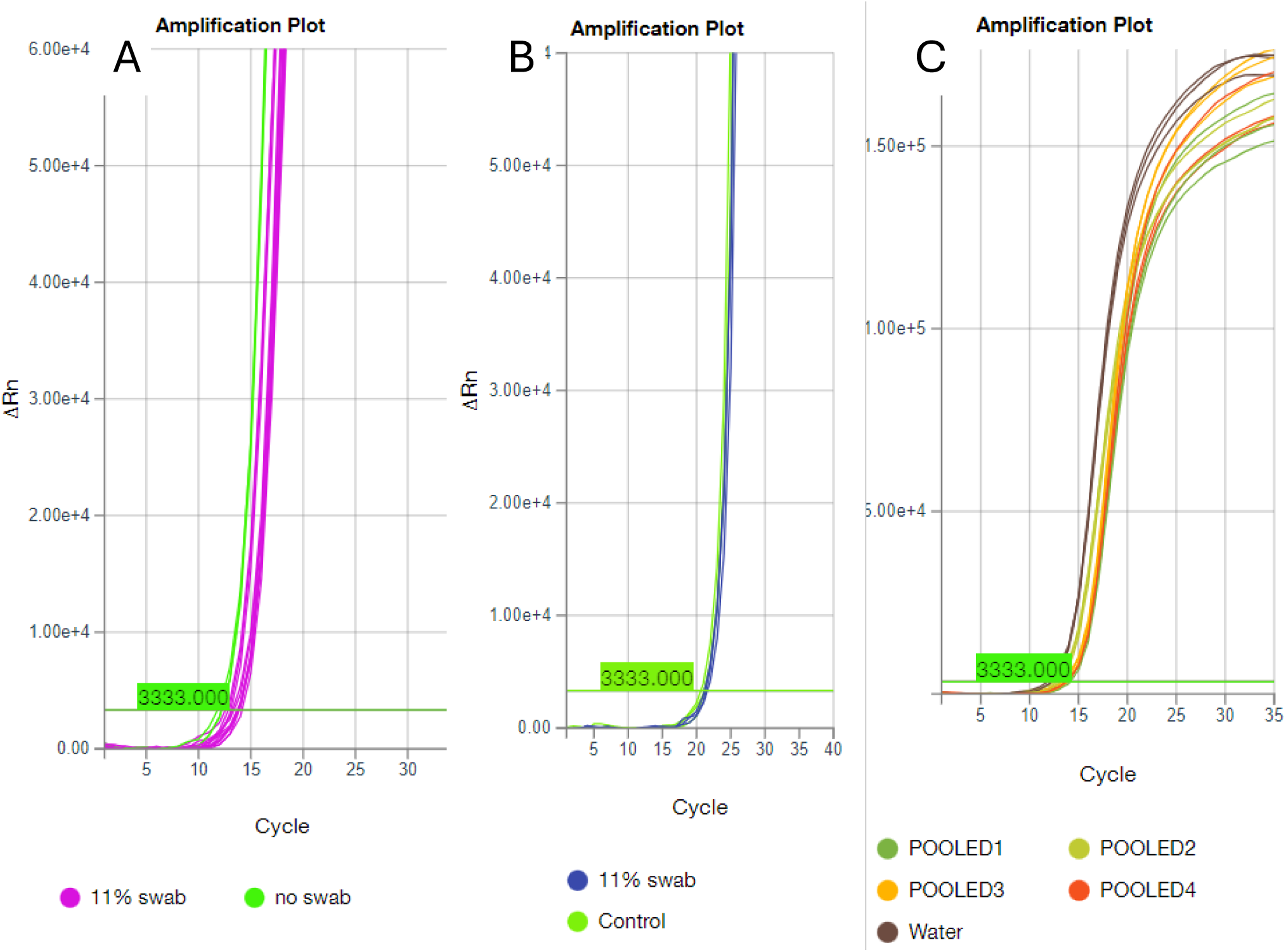
The impact of the presence of cloacal swab eluates on the AIV assay when added at 11% by volume into a reaction containing 15 IVT targets of sequence FP from table 1. a) 33 different cloacal swabs compared to a water triplicate control; b) The most inhibitory sample identified in an initial screen of 60 cloacal swab samples, tested follow process optimization; c) Demonstrating the potential use of pooled swab samples for AIV testing, five individual swabs were eluted into the same 1ml of water and used at 11% across four separate pools.

To enable more efficient testing of samples where mortality events occur or where morbidity is high, pooled sampling was assessed. For conventional diagnostic testing, sampling requires that cloacal and oropharyngeal swabs are taken from 20 individual birds where mortality is high and 60 cloacal and oropharyngeal swabs where disease presentation is nuanced. As such, with the enhanced sensitivity of the novel assays, an assessment of pooling of sample material was also advantageous. From a crude assessment, it was possible to combine swabs into the same 1ml volume of water, without any reduction in the sensitivity of the assay (**Figure 3c**), meaning that assays could practically tolerate more than the tested 11% by volume of a single swab as the inhibitors in a pooled sample would be higher.

To generate further data in support of this approach, unextracted inactivated viral antigens were spiked into cloacal and oropharyngeal swabs collected from uninfected birds, to generate synthetic swab samples. All samples were correctly identified, confirming that the assay could amplify viral products successfully in the presence of the crude samples and without the need for extraction. A demonstration that the cloacal and oropharyngeal swabs could be combined into a single 1ml reaction volume of water and then added to the reaction at a final volume of 16.6% was also completed (**Figure 4**)

**Figure 4.**
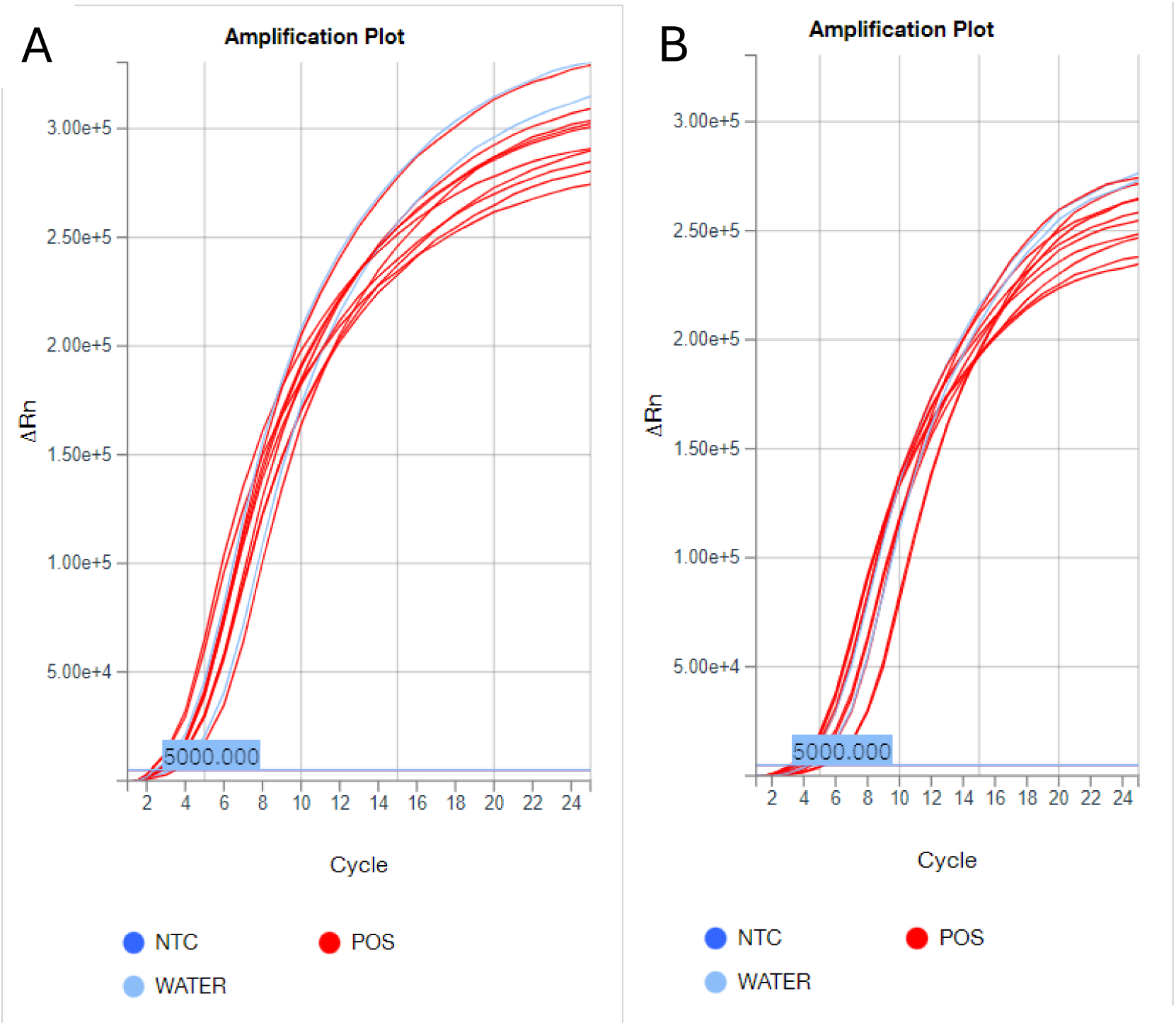
Using the inactivated antigens a) CA and b) TE (table 1), reactions were spiked with CP and OP combined swab eluate. Eluate was made by placing both a CP and OP swab into 1ml of water and using these at 16.6% by reaction volume.

We further demonstrated that the NDV assay could also be undertaken directly from unprocessed swab samples by amplifying genomic material from the NDV Ulster strain from cloacal and oropharyngeal swabs in the absence of nucleic acid extraction (**Figure 5**).

**Figure 5.**
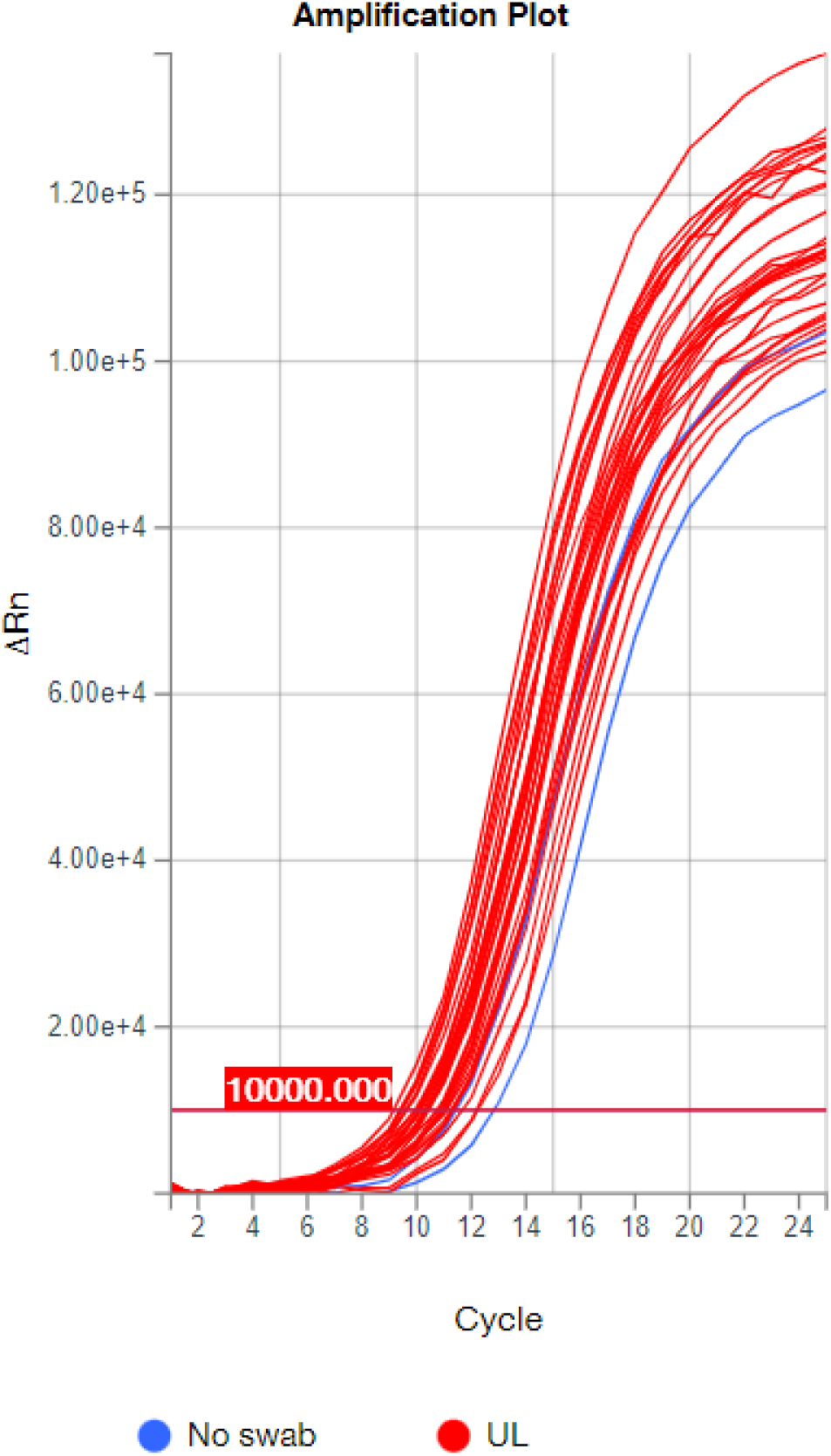
Using the UL inactivated antigen (table 1) reactions were spiked with CP or OP eluate. Data shows the impact of 14 OP and 16 CP randomly selected swabs on the performance of the NDV assay.

Importantly, whilst the current study simply supports proof of concept for these assays in a laboratory setting, the extraction-free reactions are suitable for in-field, point of need testing. This is undertaken on an innovative portable RT-PCR analysis device called CENOS with a larger reaction volume than that used in standard laboratory RRT-PCR instrumentation, to maximize the crude sample input volume. This patent-referenced technology (WO2022/195289) uses carbon-filled polypropylene to provide high thermal conductivity and low thermal mass supporting rapid thermal response and efficient heat transfer. CENOS has a biosecure consumable format that supports sealed, single-use operation and reduces contamination risk. High surface-area-to-volume geometry promotes reaction-volume temperature homogeneity and improved thermal uniformity supports reduced non-specific amplification and improved analytical sensitivity. Within the CENOS, thermal ramp performance exceeds 5°C per second, which enables rapid PCR cycling and shorter assay times. Multiplex optical detection uses an integrated micro-spectrometer for multi-channel fluorescence readout and differential panel analysis. Automated software analysis provides amplification curve calling and result interpretation for non-technical users. The closed tube assays require maximally two fixed volume liquid transfers when the reagent is supplied cold-chain free and the results calling is automated.

For testing on the CENOS, 90ul reaction volumes were used, into which the intended swab input was 15ul (16.6% v/v) per reaction. Several of the spiked swab samples were run through the CENOS device to generate initial proof of concept data (**Figure 6**).

**Figure 6.**
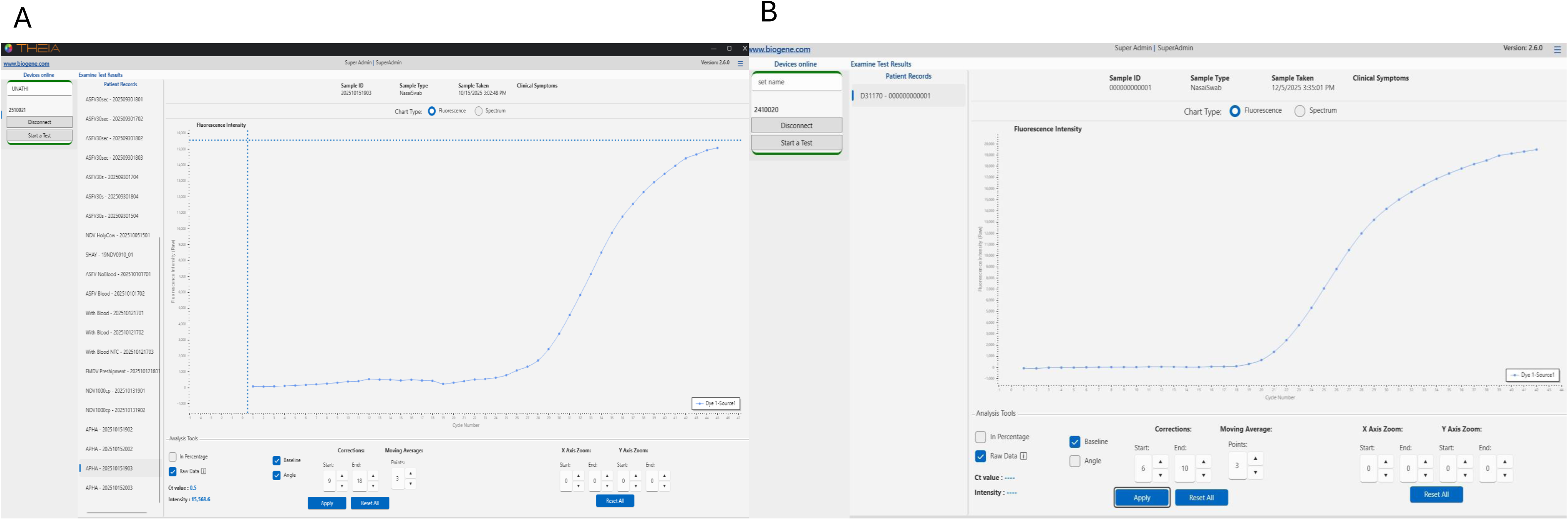
Nelson picture of cenos

As the underlying assay could detect single figure copy numbers and the swab eluate did not appear to impact the LLOD, a lower limit of detection of below 1,000 virions/ml could be envisaged. This would be more than sufficient for detecting infection in birds at a premises under suspicion of NAD as a sample set taken from a selection of birds that have either died or are exhibiting clinical disease would be collected.

Finally, to assess specificity, the assay was run using swab eluates spiked with high titres of viruses representing a panel of unrelated poultry viral pathogens including avian reovirus (AR), infectious bronchitis virus (IBV D1468, IBV 793/B), duck viral enterovirus (DVE) and duck adenovirus 1 (EDS) (Table 2). No cross-reactivity was observed for either assay with these viruses, underlining the specificity of the assay for AIV and NDV (**Figure 7**).

**Figure 7.**
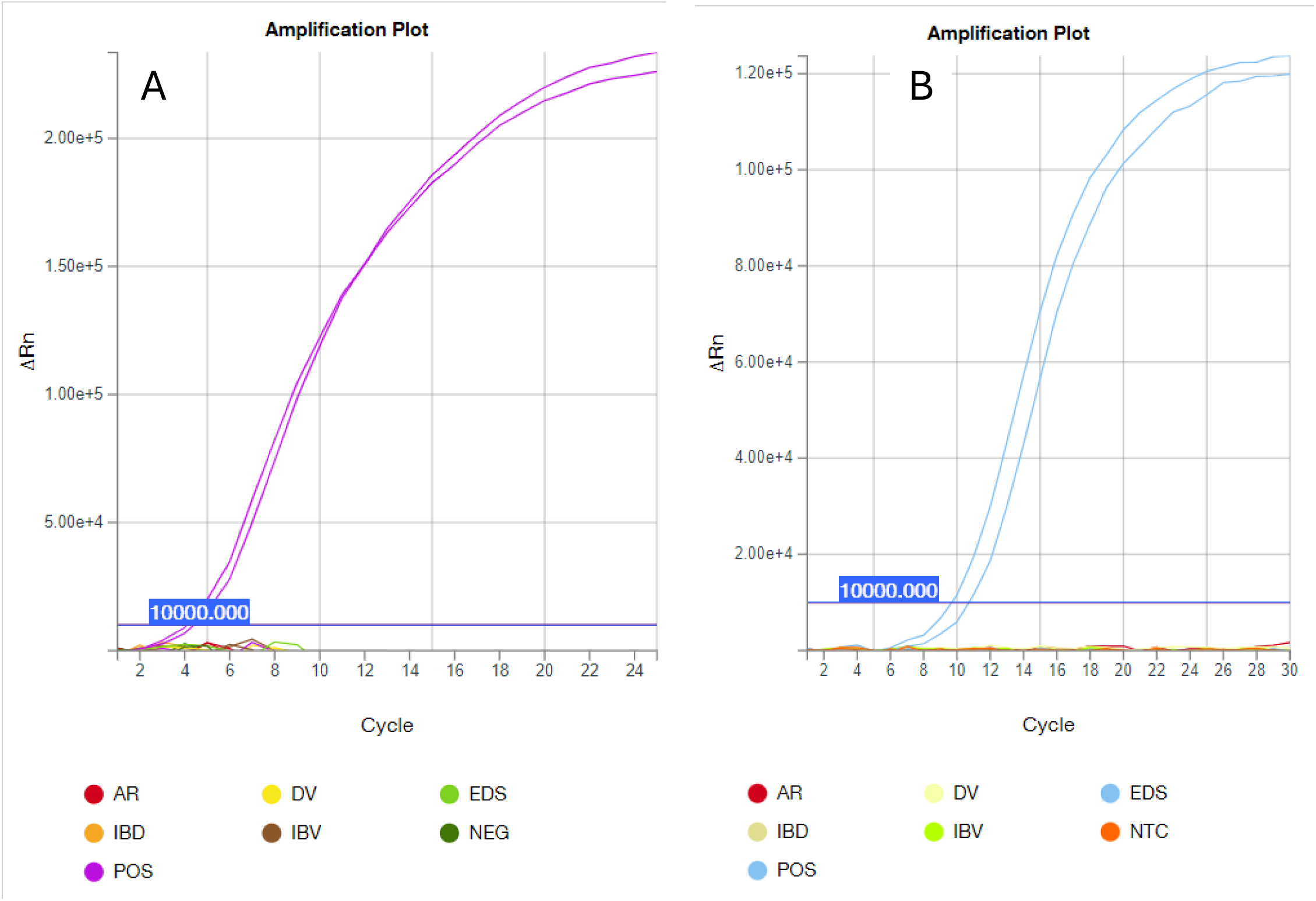
Demonstrating direct detection of AIV CA inactivated antigen template (A) and NDV UL inactivated antigen template (B) on the CENOS platform with the relevant assay. The crude sample input was 16.6% CP swab in a 90ul reaction with the following thermal cycling conditions: 94^0^C for 10s (viral lysis), 5 cycles of 69^0^C for 60s and 92^0^C for 5s (high temperature cyclical RT), 20 cycles of 68^0^C for 15s and 89^0^C for 12s, 25 cycles of 66^0^C for 10s.

**Figure 8.** The AIV assay (A) and NDV assay (B), were tested against high titres of a number of UK endemic poultry viral pathogens (Table 2) to ensure that the assays were specific for NADs and didn’t cross react with endemic pathogens. The positive samples (POS) used unextracted isolates (AIV H5N1, NDV Ulster diluted 1/10000 and added at 5ul into a 25ul reaction volume) and the other antigens tested were as outlined in table 2.

**Table 2:** Endemic viruses tested to determine assay specificity.

| Sample type | Virus / subtype | Abbreviation |
| --- | --- | --- |
| Inactivated antigen | Avian Reovirus | AR |
| Inactivated antigen | Infectious bronchitis virus (D1466) | IBV-D |
| Inactivated antigen | Duck viral enteritis | DV |
| Inactivated antigen | Infectious bronchitis virus (793/b) | IBV |
| Inactivated antigen | Egg drop syndrome virus 76 | EDS |

## Discussion

The requirement for a rapid, highly specific and highly sensitive assay that detects notifiable avian disease in a field setting is the holy grail of statutory disease diagnostics. Current lateral flow devices lack sensitivity and specificity for deployment in situations where the detection of exotic notifiable diseases is a legal requirement with profound implications associated with positive detections. The assays developed here, and their assessment against different targets, various matrices and sensitivity have generated strong proof-of-concept data to support a larger validation study for these assays employing authentic clinical outbreak samples and with direct comparison with established reference diagnostic methods. The novel assays demonstrated compatibility with higher throughput testing workflows and proved functional when applied to pooled samples as well as to concurrently collected cloacal and oropharyngeal swabs.

These two assays have demonstrated sensitivity and specificity characteristics consistent with the requirements of frontline NAD diagnostic testing when assessed against a range of currently circulating AIV genotypes and APMV-1 strains. Further, specificity was addressed through assessment against several non-notifiable avian viral diseases that are considered endemic with none being detected, demonstrating the assays specificity for NADs. Critically, the ability of these assays to detect primary targets without a reduction in sensitivity, but in the absence of undertaking RNA extraction means that the time to diagnosis is significantly reduced. Standard RT-PCR reactions require a reverse transcriptase stage of up to 30 minutes and a PCR cycling programme that often takes more than 2 hours to complete. In contrast, this extraction-free approach enables the time to diagnosis to be significantly reduced to a total of 40 minutes. Furthermore, work is currently underway to multiplex the AIV and NDV assays enabling a rapid NAD screening assay within 40 minutes. This would revolutionise testing platforms and expedite downstream activities regarding implementation of flock culling and other epidemiological requirements where NAD is detected.

Alongside the further development of the NAD specific assay, plans are underway to expand these initial assays to develop further novel approaches using the same chemistry to detect H5 and H7 viral RNA sequences, enabling the benefits of this approach to be extended to AIV subtyping. Plans are also underway to evaluate performance across a broader array of specimen types, particularly as preliminary data indicate that the current reagent system can support amplification from tissue samples.

Finally, although currently being deployed within a laboratory setting, we aim to assess the feasibility of deploying the assay outside a dedicated molecular laboratory by simplifying workflow steps, automating result interpretation, and addressing current constraints associated with handling swab material in a containment setting. This will be undertaken using the CENOS instrument which is designed for mobile, rapid and automated nucleic acid detection. The instrument performs closed-tube molecular detection direct from crude samples, with automated results calling. Reagents can be supplied cold-chain free, which enables broader long-term goals for field diagnostics. As the approach is PCR based and the device uses spectrometer-based optics, it should be possible to develop new assays for other important targets and to investigate differential diagnostic approaches. As stated, the workflow performance supports an approximately 40-minute time to result with minimal operator training. Further development of such approaches would facilitate more rapid field-based testing and potentially reduce the need to submit samples to centralized laboratory facilities.

## Acknowledgements

ACB, JJ, SMR, JT were part funded by the UK Department for the Environment, Food and Rural Affairs (Defra) and the devolved Scottish and Welsh governments under grants SE2228 and CR, ACB, JJ and SMR under grant SE2227. ACB, SMR, JT, NN and DE were funded under the Innovate UK programmes **10090901** and **10085550**. ACB and JJ were also part funded by the Biotechnology and Biological Sciences Research Council (BBSRC) and Department for Environment, Food and Rural Affairs (Defra, UK) research initiative ‘FluTrailMap’ [grant number BB/Y007271/1].

## Authorship statement

The authors declare no conflict of interest.

## Supplementary figure legends

**Supplementary figure 1.**
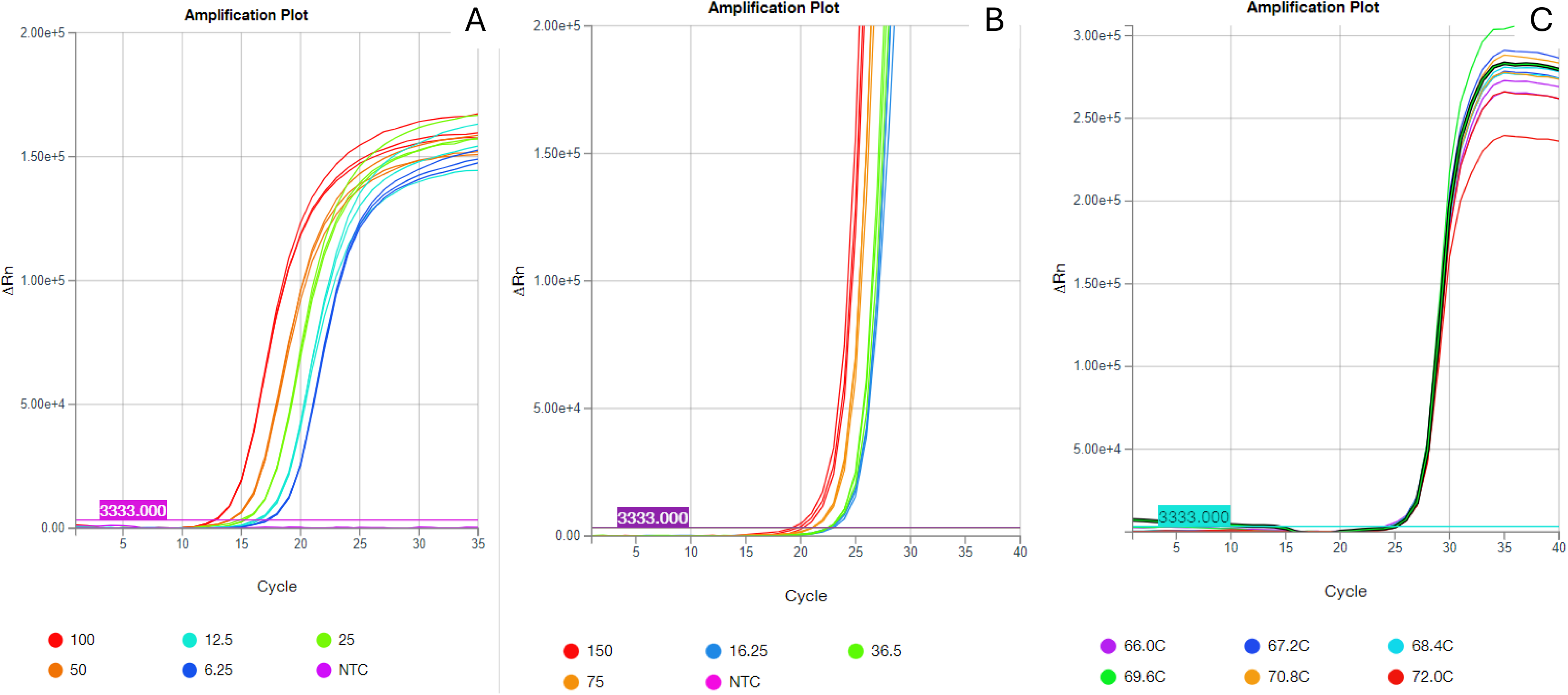
A – Showing a dilution of template near the LLOD of the AIV assay, from 100 down to 6.25 IVT targets versus NTC B – Showing a dilution of template near the LLOD of the NDV assay, from 150 down to 16.25 IVT targets versus NTC C - impact of reverse transcription temperature on the AIV assay, four 90 second reverse transcription steps were performed against a temperature gradient ranging from 66^°^C to 72^°^C

